# Reducing expression of dynamin-related protein 1 increases radiation sensitivity of glioblastoma cells

**DOI:** 10.1101/688861

**Authors:** Wen-Yu Cheng, Kuan-Chih Chow, Ming-Tsang Chiao, Yi-Chin Yang, Chiung-Chyi Shen

## Abstract

**Background:** Dynamin-related protein 1 (DRP1) is a GTPase involved in mitochondrial fission, mitochondrial protein imports, and drug sensitivity, suggesting an association with cancer progression. This study is to evaluate the prognostic significance of DRP1 in glioblastoma multiforme (GBM).

**Material and Methods:** DRP1 expression was measured by immunohistochemistry and Western blotting. Correlations between DRP1 expression and clinicopathological parameters were analyzed by statistical analysis. Differences in survival were compared by a log-rank test.

**Results:** DRP1 expression was detected in 87.2% (41/47) patients with GBM. Patients with higher DRP1 levels had worse survival (*p* = 0.0398). *In vitro*, silencing of DRP1 reduced cell proliferation, metastatic potential, and radiation resistance. The addition of shikonin inhibited DRP1 expression and increased drug uptake. Moreover, shikonin reduced the nuclear entry of DNA repair-associated enzymes and increased radiation sensitivity, suggesting that to reduce DRP1 expression could inhibit DNA repair and increase the radiation sensitivity of GBM cells.

**Conclusion:** Our results indicate that DRP1 overexpression is a prospective radio-resistant phenotype in GBM. Therefore, DRP1 could be a potential target for improving the effectiveness of radiation therapy.

Abbreviations used are: ATAD3A, the ATPase family, AAA domain containing 3A; CIM, confocal immunofluorescence microscopy; DRP1, dynamin-related protein 1; ER, endoplasmic reticulum; GBM, glioblastoma multiforme; hHR23A, human homolog of yeast Rad23 protein A; IDH1, isocitrate dehydrogenase 1; MAM, mitochondria-associated membrane; MGMT, O^6^-methylguanine-DNA-methyltransferase; SAHA, suberoylanilide hydroxamic acid (vorinostat); TMZ, temozolomide

**Translational Relevance:** This study shows that dynamin-related protein 1 (DRP1), an essential 80-kDa GTPase, which is involved in mitochondrial fission, and mitochondrial protein imports, is highly expressed in glioblastoma multiforme (GBM). Moreover, we demonstrate that DRP1 expression is closely associated with radiation sensitivity, cancer progression, and patients’ cumulative survival. *In vitro*, inhibition of DRP1 expression reduced the nuclear entry of DNA repair-associated enzymes, such as ATM, but increased radiation sensitivity and nuclear drug uptakes of glioblastoma cells. More importantly, the silencing of DRP1 induced cellular autophagy. These results indicate that DRP1 overexpression could be a prospective radio-resistant phenotype in GBM and a clinically important target for improving the effectiveness of radiation therapy.

## Introduction

Glioblastoma multiforme (GBM, Word Health Organization grade IV glioma) is the most aggressive brain tumor and thus has the worst prognosis. Most of the patients (~70%) die within two years following diagnosis. Proper radiation therapy with the pre-radiation intake of an alkylating agent, temozolomide (TMZ), has improved treatment efficacy. However, the effects have been limited [1].

Advances in molecular biology have suggested that gain of oncogene function (e.g., *N-ras*, human <u>e</u>pidermal <u>g</u>rowth <u>f</u>actor <u>r</u>eceptor [EGFR]-1 [HER-1, also known as v-ErbB-2 avian <u>e</u>rythro<u>b</u>lastic leukaemia viral oncogene homolog 1, *erbB-1*] and isoforms 1 and 2 of citrate dehydrogenase [IDH1/2]) [2, 3], as well as loss of tumour suppressor genes (e.g., p53, RB1, O^6^-<u>m</u>ethyl<u>g</u>uanine-DNA-<u>m</u>ethyl<u>t</u>ransferase [MGMT], and phosphatase and <u>ten</u>sin homolog [pTEN]) [4–7], are frequently associated with GBM. Although the oncogenic consequence is yet to be determined, risk factors in lifestyle (e.g., smoking, drinking habits and compulsive use of wireless phones) and environment (e.g., exposure to ionizing radiation and chemicals) have been implicated in the cumulative multigene alterations, which can then activate oncogene expression, induce aberrant cell growth and accelerate carcinogenic changes [8–12]. EGFR expression in GBM had attracted several provisional clinical trials targeted at EGFR-phosphatidylinositol 3-kinase (PI3K)-Akt/protein kinase B (PKB) and mammalian target of rapamycin (mTOR) signaling pathways as well as several related passages [13–15]. The preliminary results were promising; however, improvement of treatment efficacy and patient’s survival were not as evident.

Hypoxia was recently shown to be an important factor for the increase of GBM resistance simply by inducing autophagy[16]. Biochemically, hypoxia not only activated nuclear translocation of apoptosis-related mitochondrial protein, BCL-2 nineteen kilo-Dalton interacting protein 3 (BNIP3) [17, 18], but also elevated synthesis of α-ketoglutarate and 2-hydroxyglutarate by IDH to increase chromatin epigenetic modification [19, 20], as well as resistance to treatment of TMZ and radiation [1, 7, 21]. Hypoxia also induced nuclear translocation of dynamin-related protein 1 (DRP1), which was associated with DNA repair-related protein, human homolog of yeast Rad23 protein A (hHR23A), by which the DRP1 could, on one hand, protect nucleoli and, on the other hand, increase DNA repair as well as cisplatin resistance of cancer cells [22, 23].

DRP1 is an 80-kDa GTPase, which mediates budding and scission of a variety of transport vesicles and organelles [22, 24, 25], including mitochondria [26]. A number of anticancer drugs, e.g., epipodophyllotoxins and cisplatin, induce mitochondrial fragmentation, a phenomenon that is closely associated with apoptosis and chemotherapeutic cytotoxicity [27]. A better understanding of DRP1 and the enzyme effect on drug activity could, therefore, provide more valuable information to improve disease management. In addition, these chemotherapeutic agents might become vital probes for studying the essential function as well as the regulation mechanism of DRP1 and other fusion/fission-related proteins in the intracellular material trafficking and organelle damage [22–25]. However, DRP1 has not been studied in the GBM.

In this study, we used immunohistochemistry and Western blotting to determine DRP1 expression in GBM. We then evaluated the prognostic significance of DRP1 expression in GBM patients. Moreover, we investigated the effect of shikonin and suberoylanilide hydroxamic acid (SAHA, vorinostat), a histone deacetylase (HDAC) inhibitor, on DRP1 expression as well as radiation sensitivity *in vitro*.

## Materials and methods

### 1. Tissue specimens and immunohistochemical detection of DRP1 expression

From January 2008 to August 2012, tissue specimens were collected from 47 patients with newly diagnosed glioblastoma multiforme (GBM). The protocol of the study, including tissue specimen collection, pathology evaluation, the methylation status of O(6)-methylguanine-DNA methyltransferase (MGMT) promoter and survival assessment, was approved by the Medical Ethics Committee of Taichung Veterans General Hospital. Tissue microarrays of 35 American GBM samples (GL806, US Biomax, Inc., Rockville, MD, USA) were used to compare DRP1 expression between Taiwanese and American patients. Immunohistological staining was performed on formalin-fixed sections using an LSAB method (DAKO, Carpenteria, CA). The chromogenic reaction was visualized by peroxidase-conjugated streptavidin and aminoethyl carbazole (Sigma, St. Louis, MO) ([14, 22, 24, 25, 28]. Slides were evaluated by at least two independent pathologists without knowledge of the patient’s clinicopathological background. An immune-reaction scoring system was used for scoring [29]. A specimen was considered having strong signals when more than 50% of cancer cells were positively stained; intermediate signals, if 25-50% cells stained positive; weak, if the positive cells were between 10 and 25%; and negative, if less than 10% cells were stained. Cases with strong and intermediate signals (≥ 25% cells positive) were classified as DRP1^+^, those with weak or negative DRP1 signals were classified as DRP1^−^.

### 2. Cell culture and alteration of DRP1 expression using lentivirus-carrying shRNA or ectopic plasmid

Human glioblastoma multiforme cell line, U87MG, and T98G were obtained from ATCC (Manassas, VA, USA) and grown in Dulbecco modified Eagle medium (DMEM) supplemented with 10% fetal bovine serum (FBS), 4 mM glutamine, 100 U/ml penicillin and 100 μg/ml streptomycin. The cells were routinely tested and authenticated using a PromegaGenePrint^®^ 10 system for human cell line DNA typing (Mission Biotech, Taipei, Taiwan). The cells were grown to 80% confluence on the day of infection. Lentivirus carrying DRP1 shRNA was prepared using a three-plasmid transfection method [30]. The product lentivirus was used to infect T98G cells, and cells with DRP1 gene knockdown (DRP1^KD^) were selected using 1 μg/ml puromycin.

### 3. Western blotting analysis

Purified shikonin (≥98%) was purchased from Sigma-Aldrich (San Louis, Mo). Western blotting analysis has been described previously [14, 22, 24, 25, 31]. Briefly, 30 μg of total cell lysate was separated on a 10% polyacrylamide gel with a 4.5% stacking gel. After electrophoresis, proteins were transferred to a nitrocellulose membrane. The membrane was probed with specific antibodies. The protein was visualized by exposing the membrane to an X-Omat film with enhanced chemiluminescence reagent (Merck, Darmstadt, Germany). The respective primary antibodies were mouse anti-DRP1, and mouse anti-β-actin. Mouse monoclonal antibodies to DRP1 were home-made and had been characterized [22]. The digital images on X-Omat film were processed in Adobe Photoshop 7.0 (http://www.adobe.com/). The results were analyzed and quantified by the software, image-J (NIH, Bethesda, MD).

### 4. Confocal immunofluorescence microscopy

The method for immunofluorescence confocal microscopy had been described previously [22]; [24, 25]. Briefly, the cells on slides were fixed with 4% paraformaldehyde for 15 min at room temperature and permeabilized with 0.1% Triton X-100 prior to staining with mouse anti-DRP1. After washing off of the primary antibodies, slides were incubated with Alexa 488-conjugated goat anti-rabbit IgG (Invitrogen, Grand Island, NY). The nuclei were stained with 4’, 6-Diamidino-2-phenylindole (DAPI) and the slides were examined under a laser confocal microscope (Olympus FV-1000, Tokyo, Japan). Images of the cells were analyzed by FV10-ASW 3.0 software (Tokyo, Japan).

### 5. Colony formation assay

T98G, T98G-DRP1^KD^, GBM stem cells (GSC), and GSC-DRP1^KD^ cells were separately treated with 3, 6, or 12 Greys (Gy) of radiation (Varian 21EX linear accelerator, Varian Oncology Systems, Palo Alto, CA). GSC was prepared according to the previously described protocol [32]. After radiation, the attached cells were detached by treatment with trypsin and reseeded at 100, 500, 2,000, and 5,000 cells/well of culture plate, respectively. The cells were incubated at 37°C for 10 days, visible colonies that contained more than 50 cells were counted and the plating efficiency was determined. Semi-log graph of the cell survival fractions (ratio of colonies formed by irradiated cells to colonies formed by control cells) against radiation dosage was plotted.

### 6. Drug-sensitivity assay

Drug-sensitivity was measured by a WST-1 assay [33]. Cells were seeded at 100, 1,000, and 5,000 cells/96-well plates 18 hours prior to drug challenge. Cells were pulse-treated with 4 μM of daunorubicin for 2 hours. The negative control cells were treated with the solvent for the drug. Total survival of the cells was determined 72 hours following drug challenge, and percent survival was estimated by dividing optical absorbance resulted from each experiment group with that of the control group. Each experiment was done in triplicates, and the optical absorbance was measured by the coloration of the reacted substrate, WST-1 (BioVision, Mountain View, CA), which was catalyzed by mitochondrial dehydrogenases.

### 7. Statistical analysis

Overall survival (OS) was the time from the date of diagnosis to the date of death. Survival curves were plotted using the Kaplan-Meier estimator [34] and the statistical difference in survival between the different groups was compared by a log-rank test[35]. Statistical tests were two-sided, and *p* < 0.05 was considered significant. The *t*-test was utilized to compare the numerical difference of clinical parameters. Differences in patients’ performance status, tumor location, and surgical resection status were assessed by *χ*-square or Fisher’s exact test. Analyses of the data were performed using SPSS 10.3 software (Chicago, IL).

## Results

### 1. Overexpression of DRP1 in GBM specimens as determined by immunohistochemistry and Western blotting analysis

From January 2008 to August 2012, 47 GBM patients who had undergone standard surgery and palliative radiation therapy with daily TMZ (75 mg/m2) adjuvant monthly TMZ (150-200 mg/m2) were retrospectively enrolled in the study. The demography and treatment parameters of these patients are listed in Table 1. Identification and classification of tissue staining are described in detail in the Materials and Methods section. Using immunohistochemical staining, the expression of DRP1 was detected in 41 (87.2%) of Taiwanese GBM specimens (Figure 1A, as crimson precipitates in the cytoplasm), and some of the DRP1 was identified in the nuclei of tumor cells (Figure 1B, DRP1-positive nuclei were shown as brown precipitates in the nuclei, compared to DRP1-negative blue nuclei) in 33 (80.5%) of the 41 samples. The positive and negative staining controls were shown in Supplementary Figures S4A-S4C. DRP1 signal was detected in 32 (91.4%) of 35 American GBM patients, and nuclear DRP1 (DRP1^nuc+^) was detected in 27 (84.4%) specimens. No difference was found in DRP1 expression between the American and Taiwanese GBM patients (*p* = 0.609). The expression of the 80-kDa DRP1 in Taiwanese patients was confirmed by Western blotting (Figure 1C). Interestingly, molecular weights of the DRP1 in 7 of 12 surgical specimens were higher than the anticipated 80-kDa and three samples clearly had two protein bands, indicating that the DRP1 in biopsies could be post-translationally modified [22]. The above data using anti-DRP1 monoclonal antibody were home-made and had been proved and characterized[22].

**Figure 1.**
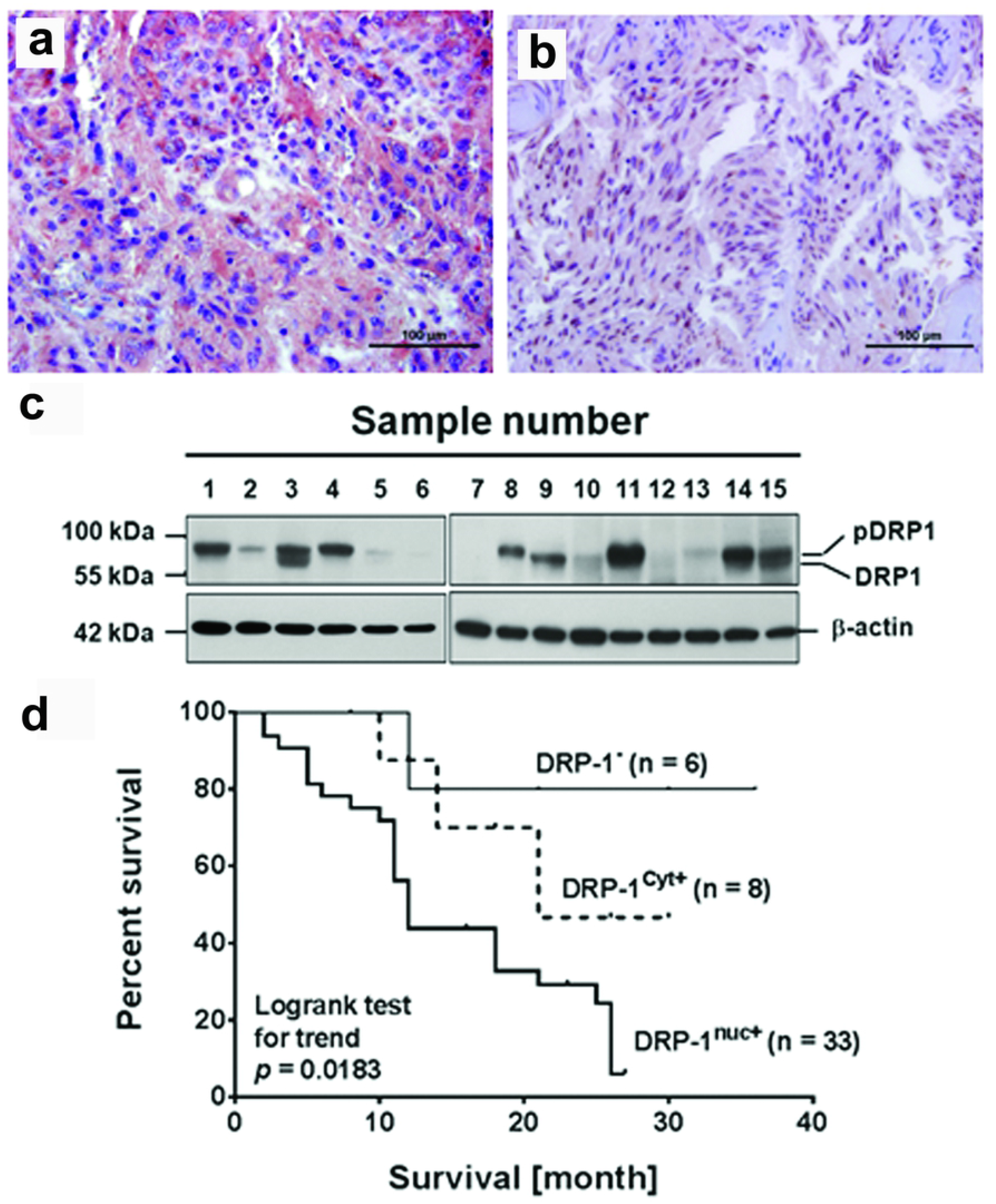
Immunohistochemical staining for the detection of DRP1 expression in GBM pathological specimens. In Taiwanese patients, (**A**) GBM cells highly expressed DRP1 (as crimson precipitates in the cytoplasm, denoted as DRP1^cyt+^) and (**B**) highly expressed nuclear DRP1 (as brown precipitates in the nuclei, denoted as DRP1^nuc+^) (original magnification × 400). The slides were counterstained with hematoxylin. (The positive and negative controls were shown in the Supplementary Figures S4A-S4C.) Scale bars are 100 μm. (**C**) Expression levels of DRP1 in surgically resected GBM specimens as determined by Western blotting. The calculated molecular weight of DRP1 was 80-kDa, and the 85-kDa protein bands are probably the phosphorylated DRP1. (**D**) Comparison of Kaplan-Meier product limit estimates of survival analysis in patients with GBM. Patients were divided into three groups, DRP1^nuc+^, DRP1^cyt+^ and DRP1^−^, according to the expression and location of DRP1. The statistical difference in survival among the three groups was compared by a log-rank test for trend. DRP1^nuc+^ patients (higher nuclear DRP1 expression) had significantly shorter OS (*p* = 0.0183). (For other survival comparisons, please check the Supplementary Figures S1A to S1E).

### 2. The impact of DRP1 overexpression on GBM patient’s prognosis

The survival of patients with low DRP1 levels was significantly better than that of patients with high DRP1 levels. The difference between cumulative overall survival (OS) [*p* = 0.0398, 95% confidential interval (CI), 1.051-8.151; Hazard ratio (HR) between DRP1^+^ and DRP1^−^ patients was 5.71) were significant (Supplementary Figures S1A & S1B). The actual 18 month OS rate of DRP1^+^ patients was 40.0%, while that of DRP1^−^ patients was 80.0%. Survival of DRP1^−^ patients was indeed better than DRP1^+^ patients. When nuclear DRP1 was used as a perspective parameter, survival of patients with nuclear DRP1 was significantly worse than that of the other two groups (Figure 1D, *p* = 0.0183, log-rank test for trend; or Supplementary Figures S1C and S1D, OS, *p* = 0.0039, and PFS, *p* < 0.0001), indicating that expression of DRP1, including nuclear DRP1, could act as a prognostic phenotype of GBM. Subgroup analyses revealed that GBM patients with DRP1 overexpression and unmethylated MGMT promoter had the worst radiation responses and survival (Supplementary Figures S1E & S1F). At the time of data analysis (patients had been routinely followed for up to 24 months), 5 (83.3%) of 6 DRP1^−^ patients were alive. Among these, four were progression-free.

### 3. Silencing of DRP1 Expression in GBM Cells decreases cell growth, and mobility, but increases radiation sensitivity

*In vitro*, protein levels of DRP1 were examined by Western blotting analysis in a mouse brain tumor cell line (H4), and two human glioma cell lines (U87MG and T98G). Al three cell lines expressed both 80- and 85-kDa proteins (Figure 2A). Identities of the immunoprecipitated proteins were determined by matrix-assisted laser desorption/ionization-time-of-flight-mass spectrometry (MALDI-TOF). Peptide mass fingerprinting of both 80-kDa and 85-kDa proteins matched to DRP1: O00429, DRP1, indicating that both 80-kDa and 85-kDa proteins were DRP1 and that the 85-kDa protein could be post-translationally modified (Supplementary Figures S2A-2D).

**Figure 2.**
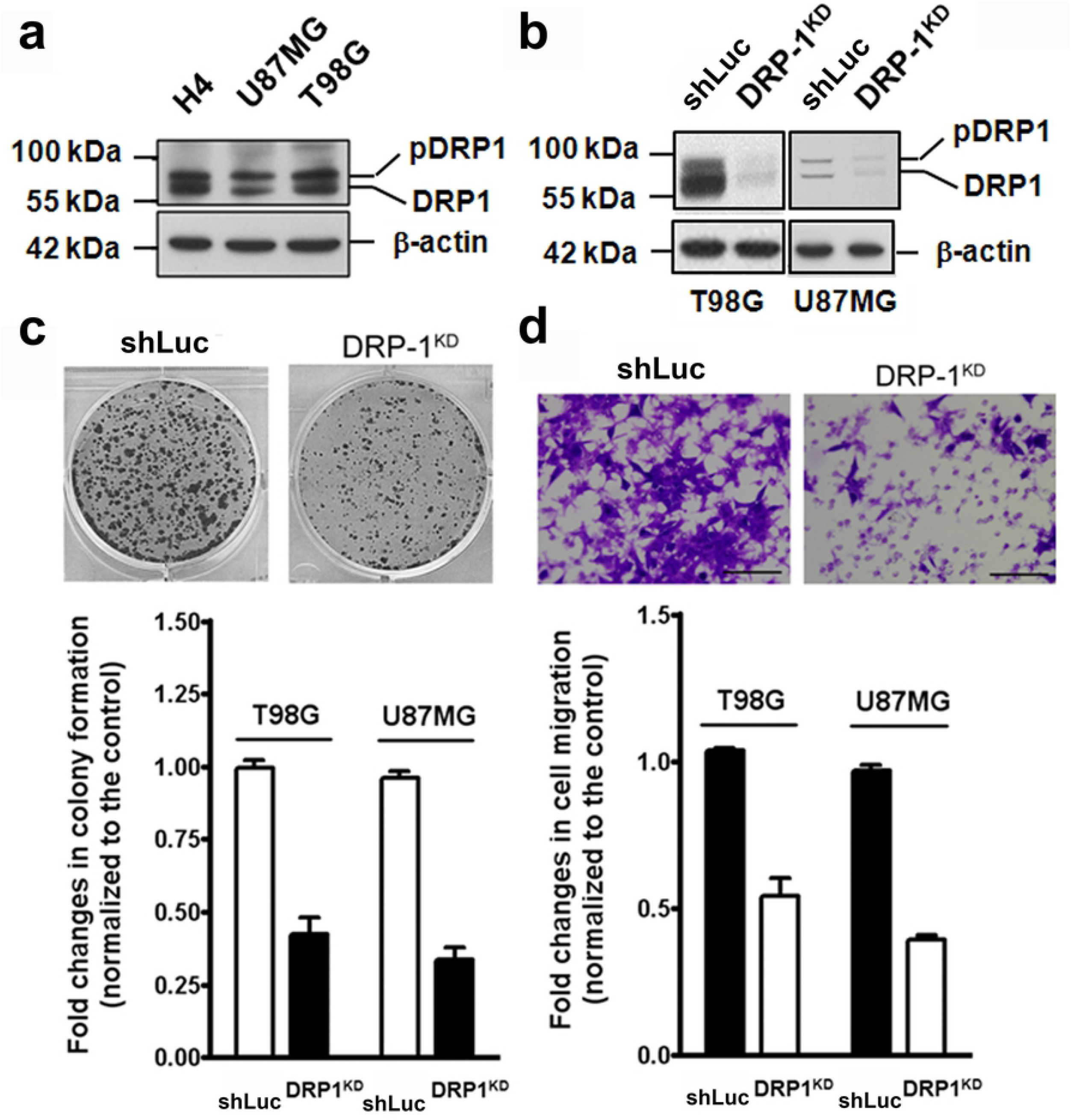
Correlation of DRP1 expression with cell growth and invasion potential of GBM cells. (**A**) DRP1 was highly expressed in mouse H4 and human T98G cells. DRP1 level in U87MG was lower than that in T98G cells. (**B**) The silencing of DRP1 expression (DRP1^KD^) reduced the DRP1 protein level (as detected by Western blotting) of the T98G and U87MG cells. (**C**) The silencing of DRP1 expression decreased proliferation capacity (as measured by colony formation and WST-1 assays). (**D**) DRP1^KD^ T98G and DRP1^KD^ U87MG cells had lower invasion potential (as measured by matrigel penetration assay) Scale bars are 250 μm. The results were repeated over three independent experiments in each case.

As noted above, both pathological and clinical studies showed that higher DRP1 expression correlated with worse prognosis in patients concurrently treated with TMZ and irradiation. We, therefore, examined the effect of DRP1 on cell proliferation and migration. *In vitro*, inhibition of DRP1 expression by using shRNA to knockdown DRP1expression (DRP1^KD^) (Figure 2B) reduced cell growth (Figure 2C) and mobility of tumor cells across matrigels (Figure 2D).

The current studies have provided that CD-133^+^ GBM stem cells retained more resistance than cancer cells to ionizing radiation.[36, 37] Silencing of DRP1, on the other hand, increased radiation sensitivity in both T98G (Figure 3A) and U87 cells (Figure 3B). The addition of TMZ only increased radiation sensitivity of DRP1^KD^ T98G cells. The decrease in radiation resistance was about 5-10 folds. Interestingly, CD-133^+^ GBM stem cells (GSC) also highly expressed DRP1, in particular, the 85-kDa protein (Figure 3C). The silencing of DRP1 expression inhibited cell growth of GSC (Figure 3D). These results confirmed our previous findings that DRP1, essential for mitochondrial protein import, was involved in cell growth and genotoxic resistance [22, 24, 25], suggesting that reducing total intracellular DRP1 expression or nuclear DRP1 levels could enhance anticancer efficacy of radiation and anticancer drug therapies.

**Figure 3.**
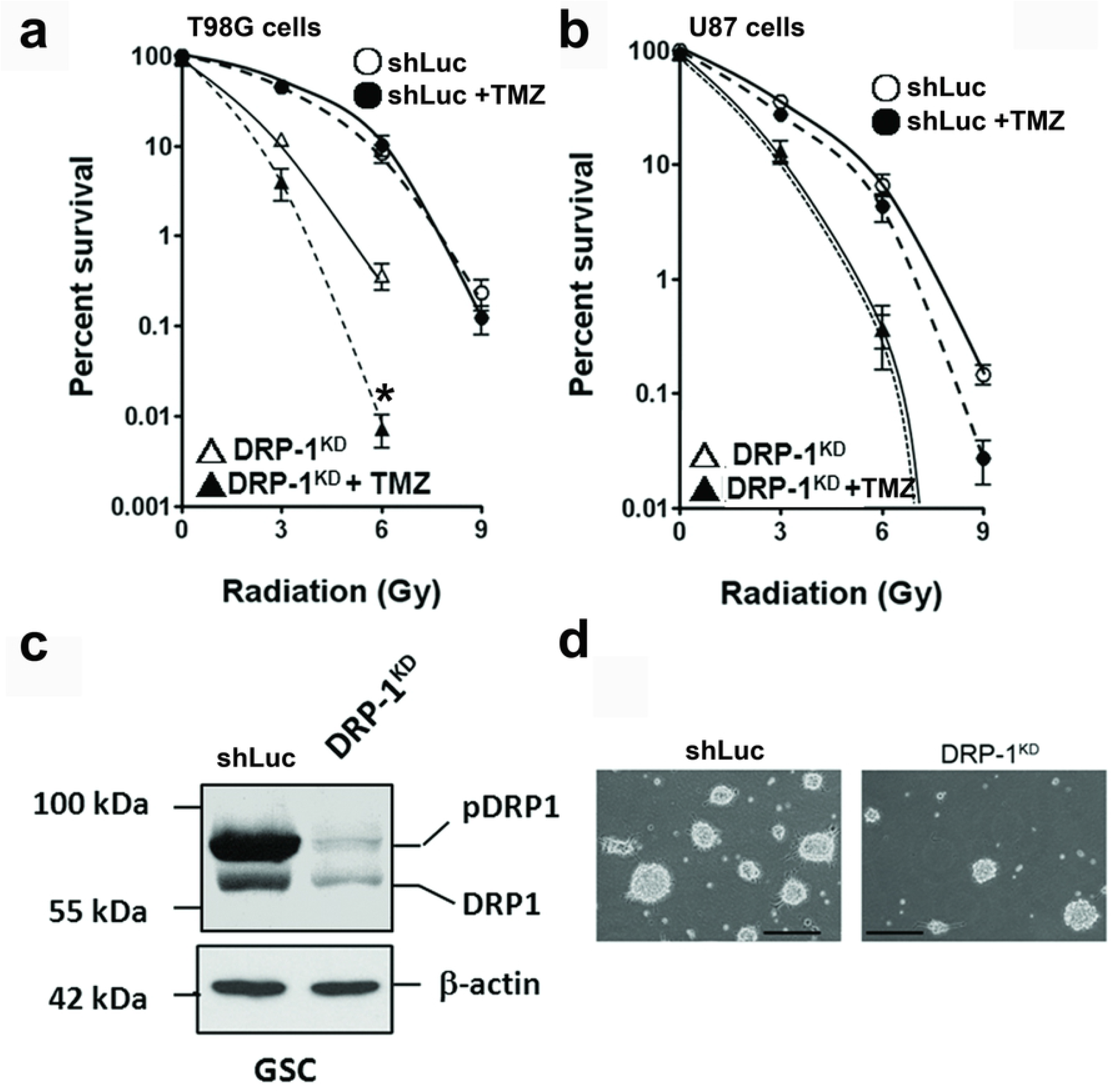
Correlation of DRP1 expression with radiation sensitivity of GBM and proliferation efficiency of GBM stem cells. (**A**) The silencing of DRP1 (DRP1^KD^) increased radiation-induced cell death (as measured by colony formation assay) in T98G cells. The addition of 50 μM TMZ did not affect radiation resistance in wild-type (●), but increased radiosensitivity of DRP1^KD^ (▴) T98G cells. ○, wild-type; △, DRP1^KD^ cells. (**B**) The silencing of DRP1 (DRP1^KD^) also increased radiosensitivity in U87MG cells. ○, wild-type; △, DRP1^KD^. TMZ reduced radiation resistance in wild-type U87MG (●), but not evidently in DRP1^KD^ (▴) cells. MGMT promoter in U87 is methylated, and that in T98G is unmethylated. Results are the means ± S.D. of three independent experiments. *, *p* <0.005 (**C**) GBM stem cells highly expressed DRP1. The silencing of DRP1 expression (DRP1^KD^) reduced the DRP1 protein level (as detected by Western blotting). (**D**) DRP1^KD^ GBM stem cells had lower proliferation ability (as measured by the formation of spheres). Scale bars are 250 μm. The results were repeated over three independent experiments in each case.

### 4. The respective effects of shikonin and SAHA on DRP1 expression and cell survival

Our previous studies showed that DRP1 was involved in an alternative mitochondrial import, and disruption of this passage induces autophagy [24, 25]. Using DRP1 as a target, we found that several Chinese medicinal herbal extracts (CMHEs) inhibited DRP1 expression, including *Astragalus, propinquus, Koelreuteria elegans, Lithospermum erythrorhizon*, and *Polygala tenuifolia* [38]. Using the web engine (http://www.google.com.tw/) to search for the major ingredients of the plants, we found that shikonin from the *L. erythrorhizon* is one of the most promising pure compounds. To evaluate the effects of shikonin and DRP1 protein on the process of autophagy and apoptosis, we analyzed the related protein expression.

As shown in Figure 4A, shikonin decreased both 80- and 85-kDa DRP1, and increased autophagic marker, LC3B-II. Using fluorescence microscopy, shikonin clearly induced the formation of autophagosomes (Figure 4B). Although SAHA did not affect DRP1 expression (Figure 4C), it clearly increased levels of poly [ADP-ribose] polymerase 1 (PARP-1), a marker of apoptosis (Figure 4D, upper panel), but did not induce cleavage of PARP-1.

**Figure 4.**
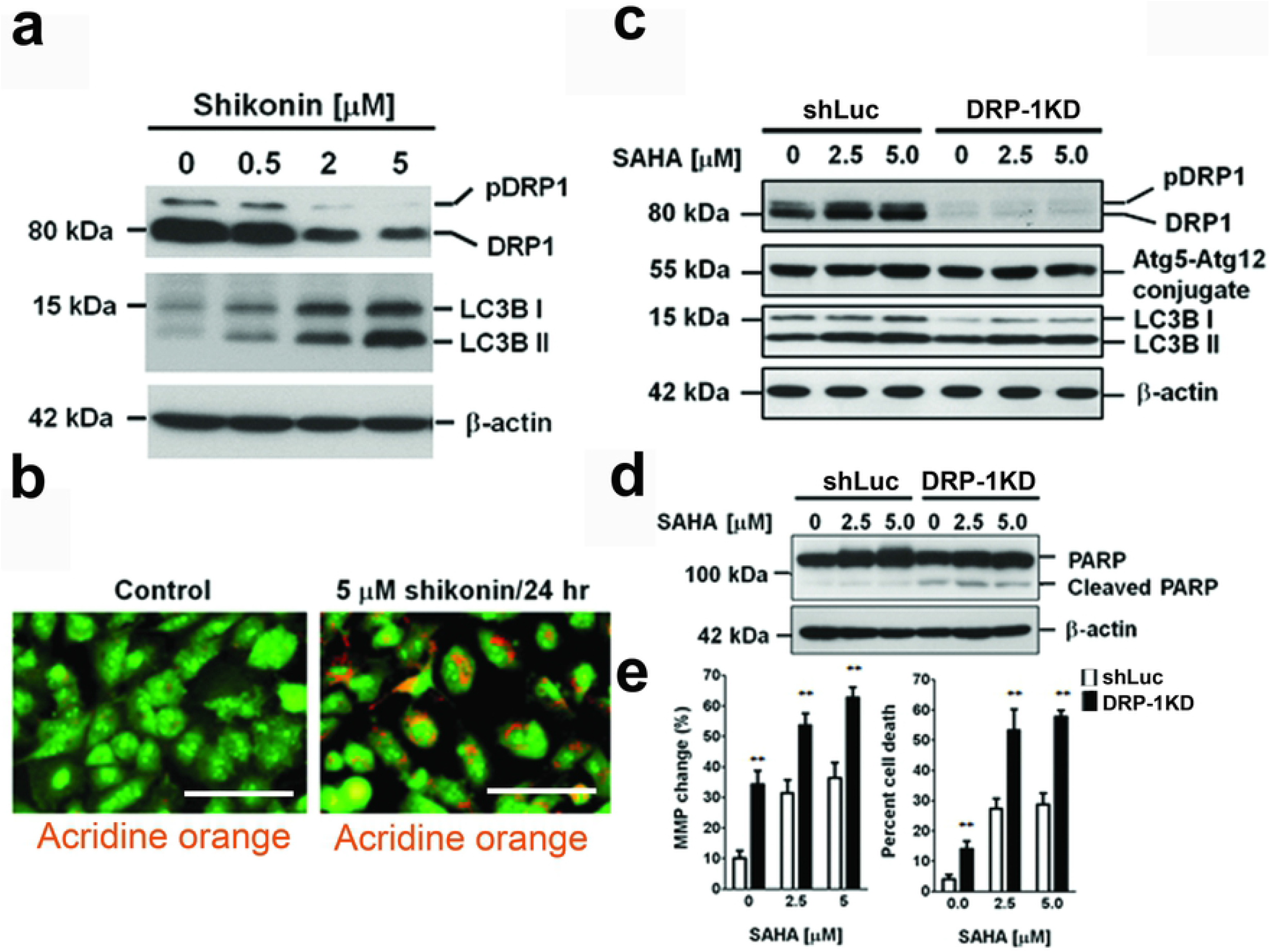
Shikonin and SAHA had different effects on gene expression of DRP1 in T98G cells. (**A**) The addition of shikonin inhibited expression of DRP1, and induced autophagy (as shown by increased levels of LC3B-II by Western blotting). (**B**) Treatment with shikonin for 24 hours induced autophagy, as confirmed by the formation of autophagosomes, which was visualized by using fluorescence microscopy to detect the colour change of acridine orange (change from colorless to yellow or orange under low pH). Scale bars are 100 μm. (**C**) The addition of SAHA did not clearly affect DRP1 expression or induce autophagy (no obvious change of autophagy markers, ATg5-Atg12 conjugates and LC3B II, was detected by Western blotting). (**D**) Treatment with SAHA for 24 hours did not readily induce cleavage of poly [ADP-ribose] polymerase 1 (PARP-1), a marker of apoptosis, but clearly increase PARP-1 levels (left side). In DRP1^KD^ T98G cells, SAHA increased PARP-1 cleavage (right side). Expression of β-actin was used as a monitoring standard for relative protein expression in the Western blotting analysis. (E) Cell death, which was measured by colony formation assay, was presented in the right panel. Changes of mitochondrial membrane potential (MMP), an indication of mitochondria depolarization, was shown in the left panel. Briefly, following SAHA treatment, T98G cells were incubated with hydrophobic fluorescent dye 3,3’-dihexyloxacarbocyanine iodide (DiOC_6_) at 37°C for 20 min prior to harvest. The collected cells were analyzed by the FACS Calibur (BD, CA, USA). Results are the means ± S.D. of three independent experiments. **, *p* <0.001

In DRP1^KD^ T98G cells, SAHA increased PARP-1 cleavage (Figure 4D, right hand side) as well as cell death (Figure 4e right-panel) and mitochondria depolarization [Figure 4e left the panel, as shown by changes of mitochondrial membrane potential (MMP)]. After SAHA treatment, T98G cells were harvested and respectively analyzed by Western blotting and flow cytometry. Expression of β-actin was used as a monitoring standard for relative protein expression in the Western blotting analysis. Results are the means ± S.D. of three independent experiments. \*\**p* <0.001

### 5. Shikonin increases nuclear levels of anticancer drugs and arrests of DNA repair-related proteins in the perinuclear MAM

Our previous studies showed that inhibiting intracellular cargo transportation-related enzymes could result in a reduction of nuclear levels of DNA repair-related proteins, such as ataxia-telangiectasia-mutated (ATM) kinase, and an increase of the cytotoxic effect of anticancer drugs and irradiation [22, 24, 25, 39]. Interestingly, using an Operatta^®^ high content imaging system (PerkinElmer, Waltham, MA) to examine the effect of shikonin on the nuclear levels of 4’,6-diamidino-2-phenylindole (DAPI) and daunorubicin, in T98G cells, we found that shikonin not only markedly increased nuclear DAPI and daunorubicin, but also significantly increased cell sensitivity to daunorubicin (Figures 5A–5C). Moreover, shikonin treatment reduced the nuclear accumulation of ATM (Figure 5D, the left panel), supporting our former results that inhibition of DRP1 expression restricted nuclear import of DNA repair-related enzymes and induced bulging of MAM (Figure 5D, right panel). Using a transmission electron microscopy, we further showed that shikonin treatment increased nuclear envelop damages (Figures 5E1 & 5E2).

**Figure 5.**
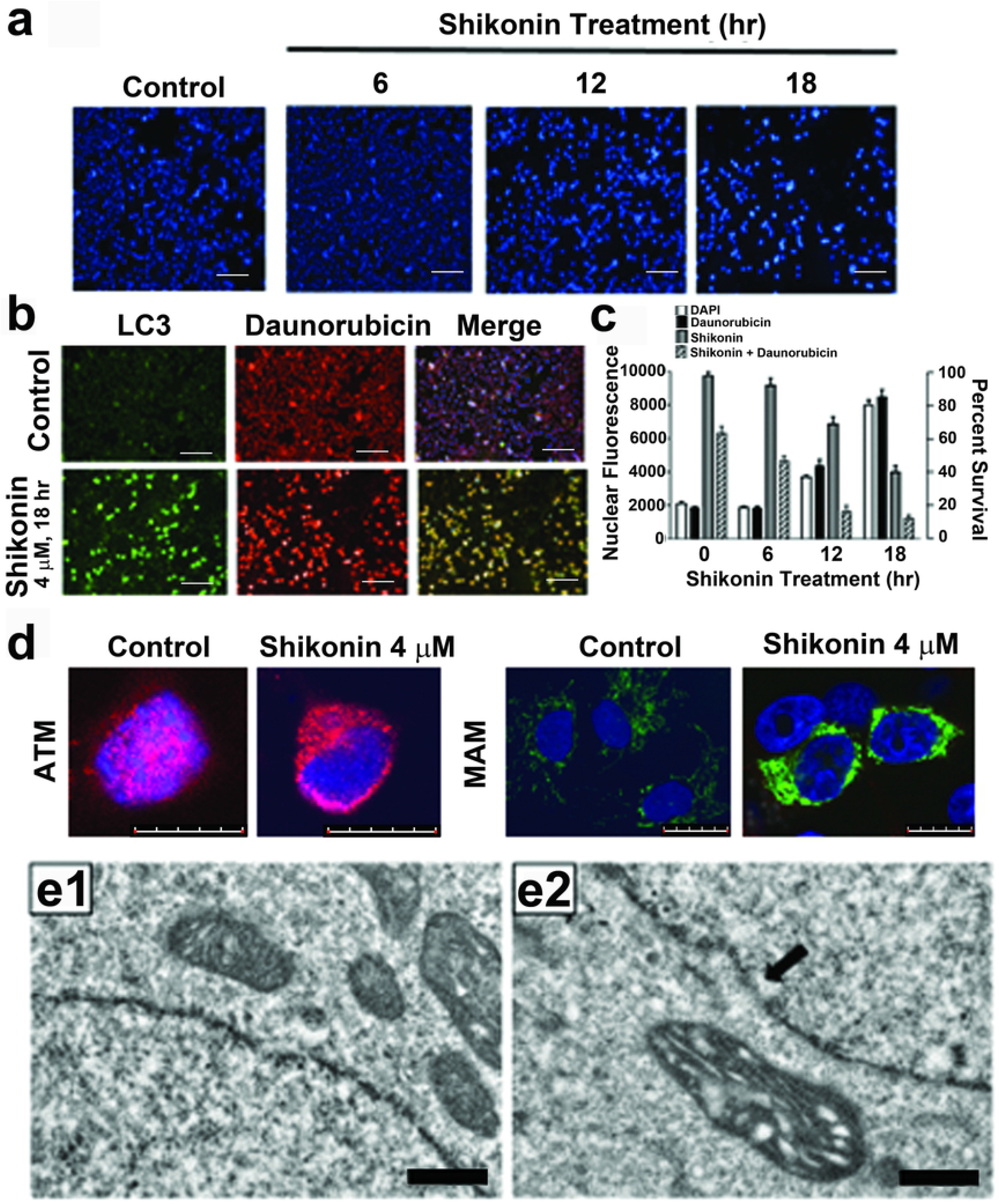
Shikonin increases nuclear levels of DAPI stain and anticancer drug, daunorubicin, but inhibits nuclear transportation of DNA repair-related protein, ATM. (**A**) 18-hr post-shikonin treatment, nuclear fluorescence of DAPI stain increased about 4 folds when T98G cells were scanned by an Operatta^®^ imaging system. Scale bars are 250 μm. (**B**) Shikonin increased expression of LC3, an autophagy marker, in T98G cells. The LC3 signals were overlapped with the fluorescence of anticancer drug daunorubicin when cells were scanned by Operatta^®^ imaging system. Scale bars are 250 μm. (**C**) A schematic composite of shikonin-treated cells. White column, cells stained with DAPI (as fluorescence control); Black column, nuclear levels of daunorubicin; Grey column, cytotoxicity to shikonin alone; Slash-line column, cytotoxicity to shikonin and 0.5 μM daunorubicin, as measured by a WST-1 assay. The results were repeated over three independent experiments in each case. (**D**) Shikonin inhibited the nuclear transportation of ATM (red fluorescence), and the proteins were accumulated in the enlarged MAM (green fluorescence), indicating that decreasing expression of DRP1 also reduced nuclear import of DNA repair-related proteins (You *et al*, 2013). Scale bars are 20 μm. The above results were repeated over three independent experiments in each case. (**E**) Comparing to the control T98G cells (**E1**) shikonin induced damage of the nuclear envelops (**E2**, arrow) when the cells were examined by a transmission electron microscopy. Scale bars are 2.5 μm.

## Discussion

Our results show that DRP1 is highly expressed in newly diagnosed GBM patients (87.2%, 41/47). Moreover, nuclear DRP1 was identified in 33 (80.5%) of DRP1-positive (DRP1^+^) pathological specimens. Using Western blotting to analyze DRP1 expression, we found that the molecular weights of DRP1 in 10 of 12 surgical samples were around 85-kDa. In spite of the number of analyzed patients are less, but it still moderated indicated that DRP1 in GBM biopsies could be post-translationally modified [16, 22]. Statistical analyses showed that patients with DRP1 overexpression or nuclear DRP1 (DRP1^nuc+^) were more resistant to radiation and hence had a higher frequency of disease relapse and worse prognosis. Subgroup analyses revealed that GBM patients with DRP1 overexpression and unmethylated MGMT promoter had the worst radiation responses and survival (Supplementary Figures S1E & S1F).

*In vitro*, DRP1 expression correlates with resistant phenotype to radiation and TMZ (T98G cells were more resistant than U87MG cells). Nonetheless, silencing of DRP1 gene increases sensitivity of both U87MG cells (with a methylated MGMT promoter) and T98G cells (with an unmethylated MGMT promoter) to radiation and TMZ, suggesting that methylation status of MGMT promoter may affect expression of DRP1, as well as that of Aldo-keto reductase (AKR) 1C1 and 1C2 [39, 40] to enforce radiation phenotype of GBM cells [22] [14, 24, 25, 31]. Binding of DRP1 to the nucleoli could further protect the rRNA-encoding region to maintain genome stability [22], and these events together could regulate cellular activity against cytotoxic agents and radiation.

Interestingly, long-term exposure of GBM cells to TMZ decreases drug sensitivity by up-regulating the expression of AKR enzymes and glucose transporter [41]. Elevation of glucose transport altered mitochondrial metabolism, while the increase of AKR enzymes deactivated TMZ and cisplatin, supporting our findings that some of AKR enzymes were localized on the mitochondria-associated membrane (MAM), the essential organelle that regulated material transports to mitochondria and nucleus [38, 39]. Both transportation passages require DRP1, ATAD3A, and mitofusin 2 (Mfn2) [25, 42]. Since shikonin inhibits DRP1 expression in T98G cells, it is reasonable to believe that intracellular materials, such as proteins and lipids which are synthesized in the endoplasmic reticulum (ER), and scheduled to be transported to mitochondria and nucleus [22, 25], will be accumulated in the MAM. The lack of timely material supply would be difficult to maintain mitochondrial integrity, which could severely diminish the mitochondrial function and change the organelle morphology (Supplementary Figures S3A-S3D).

It is worth noting that mitochondria do not synthesize phosphatidylserine (PS) *per se*. The PS is mainly synthesized in the ER and MAM, and imported to the mitochondria. *Vice versa*, the phosphatidylethanolamine (PE), the unique phospholipid that is conjugated to <u>a</u>utophagy-<u>r</u>elated <u>g</u>ene 3 (Atg3) during initiation of autophagy, is converted from the PS in the mitochondria and transported back to the ER. Interestingly, the ER also constitutes the outer part of the nuclear envelope as well. It is, therefore, reasonable to anticipate that a decrease of cytoplasmic DRP1 may concurrently damage the mitochondrial membrane and the nuclear envelope, which not only decreases general ATP supply but also reduces nuclear imports of DNA repair-related enzymes [39]. Moreover, elevated nuclear import of DRP1 could consume massive intracellular hHR23A, which would competitively diminish the nuclear import of xeroderma pigmentosum complementation group C (XPC) to delay nucleotide excision repair (NER) that was essential for maintaining genome integrity following the challenge of TMZ or cisplatin [22, 39, 41, 43].

Both TMZ and radiation induce nuclear and mitochondrial genome DNA breakage. TMZ affects mitochondrial electron transports and oxidative phosphorylation as well [44]. Radiation, on the other hand, induces the translocation of ATM, which is important for the repair of DNA breaks, to the nucleus and mitochondria [45]. ATM deficiency, either by a genetic or a biochemical method, reduces genomic DNA repair function as well as mitochondrial biogenesis and oxidative respiratory function [46]. By demonstrating that extranuclear ATM bound to ER-associated peroxisome targeting signal type 1 (PTS1) receptor (also named peroxisomal biogenesis factor 5, Pex5), Watters *et al* suggested that besides nucleus, ATM could be targeting to the MAM [47]. In a gene knockout study, Baumgart *et al* further showed that defect in the *Pex5* gene reduced peroxisomal metabolism, as well as expression and activities of mitochondrial respiration system [48]. Their results strongly suggested that MAM and its associated enzymes, in particular, the DRP1, a GTPase, played a pivotal role in allocating materials, which were essential for maintaining organelle morphology, and DNA integrity of the genome and the mitochondria. Our data supported their results and showed that reducing cytoplasmic DRP-1, either by addition of shikonin or exposure to hypoxia (Supplementary Figures S4A-S4D), might inhibit import of DNA repair-associated enzymes [39, 43] and that of mitochondrial biogenesis- and oxidative respiration-related proteins to decrease genomic and mitochondrial DNA stability, which was ultimately reflected in an increased sensitivity to drugs and radiation.

Autophagy generally regarded as a rescue response under cells including both normal cells and tumor cells, are confronted with danger, such as starvation, irradiation exposure. The inhibition of DRP1 significant increased the radiation sensitivity and repressed the autophagy response under cells faced chemo-treatment (Figures 3AB, 4C), at the same time, DRP1-KD led to an increase apoptosis response of glioblastoma cells (Figure. 4D). Obviously, the lack of DRP1 similar to the inhibition of autophagy contributed to the cell’s sensitivity to both chemo- and radio-resistance.[32, 49, 50] In spite of this study couldn’t explain clearly the played role of DRP1 in the autophagy process, while revealing evidence of light is likely to the DRP1 participated with mitochondrial DNA stability (Figure. 4e)[51, 52].

In conclusion, our results showed that DRP1 was overexpressed in GBM. The inhibition of DRP1 expression induces autophagy and enhances radiation sensitivity. This effect is specific to cancer cells, which overexpress not only DRP1 but also ATAD3A, AKR1C1, eukaryotic elongation factor (eEF2) and optic atrophy 1 (OPA1), a condition that is not detected in non-tumor counterpart [24, 31, 53]. In addition to inducing autophagy, silencing of DRP1 reduced cell growth and invasion potentials, and such features were also found in GBM stem cells. Reducing DRP1 expression augmented the cytotoxicity to SAHA (acetylation and apoptosis-inducing agent) and daunorubicin as well. Although the size of this study population was small, our data shed some light on the radio-resistant phenotype of GBM, of which DRP1 could be a potential marker; though DRP1 alone might not be an independent prognostic factor.

## Acknowledgements

This work was in part supported by the Comprehensive Academic Promotion Projects (NCHU 1025025, Ministry of Education, Taiwan) and the Ministry of Education, Taiwan, ROC, under the ATU plan, and in part by clinical research grants from Taichung Veteran General Hospital (101DHA0500377), Taichung, Taiwan; the Department of Health, Executive Yuan, Taipei, Taiwan to the China Medical University Hospital, Excellence in Cancer Research program (DOH102-TD-C-111-005), Taichung, Taiwan; and the National Science Council (NSC 102-2320-B-005-006), Taipei, Taiwan. The funders had no role in study design, data collection, and analysis, decision to publish, or preparation of the manuscript.

## Authors’ Contributions

**Conception and Design:** W Cheng, K Chow, C Shen; **Development of methodology:** K Chow; **Acquisition of data:** W Cheng, C Shen; **Analysis and interpretation of data:** W Cheng, MT Chiao, YC Yang, K Chow; **Writing and review:** W Cheng, K Chow, C Shen; **Administrative, technical or material support:** W Cheng, K Chow

